# Beyond synthetic lethality in large-scale metabolic and regulatory network models via genetic minimal intervention sets

**DOI:** 10.1101/2025.11.07.687323

**Authors:** Naroa Barrena, Carlos Rodriguez-Flores, Luis V. Valcárcel, Danel Olaverri-Mendizabal, Xabier Agirre, Felipe Prósper, Francisco J. Planes

**Affiliations:** University of Navarra, Tecnun School of Engineering, Manuel de Lardizábal 13, 20018 San Sebastián, Spain; University of Navarra, Biomedical Engineering Center, Campus Universitario 31009 Pamplona, Navarra, Spain; University of Navarra, Instituto de Ciencia de los Datos e Inteligencia Artificial (DATAI), Campus Universitario, 31080, Pamplona, Spain; Hematology-Oncology Program, CIMA, Cancer Center Clínica Universidad de Navarra (CCUN), Pamplona, Spain; CIBERONC Centro de Investigación Biomédica en Red de Cáncer, Pamplona, Spain; IdiSNA Instituto de Investigación Sanitaria de Navarra, Pamplona, Spain; Hematology and Cell Therapy Service, Cancer Center Clínica Universidad de Navarra (CCUN), Pamplona, Spain

## Abstract

**Motivation:** The integration of genome-scale metabolic and regulatory networks has received significant interest in the field of Cancer Systems Biology. However, the identification of lethal genetic interventions in these integrated models remains a substantially challenge due to the combinatorial explosion of potential solutions. To address this issue, we previously developed the genetic Minimal Cut Set (gMCS) framework, which is capable of computing synthetic lethal interactions - minimal sets of gene knockouts that are lethal for cellular proliferation- in genome-scale metabolic networks with signed directed acyclic regulatory pathways. Here, we present a novel formulation to calculate genetic Minimal Intervention Sets, gMISs, which incorporate interventions with both gene knockouts and knock-ins.

**Results:** With our gMIS approach, we assessed the landscape of lethal genetic interactions in human cells and the impact of gene knock-in perturbations, which enable us to capture for the first-time genetic interventions beyond synthetic lethality, particularly synthetic dosage lethality and tumor suppressor gene complexes. We applied the concept of synthetic dosage lethality to predict essential genes in cancer and demonstrated a significant increase in sensitivity when compared to large-scale gene knockout screen data. Moreover, we conducted a tumor suppressor analysis in cancer cell lines and identified single gene knock-in strategies that block cellular proliferation. Finally, we illustrate the ability of our gMIS approach to elucidate potential targets for cancer research with several examples in hematological malignancies

**Availability and implementation:** The Python package gMCSpy has been updated to include the functionalities to calculate gMISs. It can be accessed here: https://github.com/PlanesLab/gMCSpy.

## 1. Introduction

Synthetic lethality (SL) defines a type of genetic interaction where the co-occurrence of two (or more) genetic events results in cellular death, while the occurrence of either event on its own is compatible with cell viability (O’Neil *et al*., 2017). The concept of SL has been widely studied in the context of cancer research from very different perspectives, including large-scale genetic screens and computational methods (Wang *et al*., 2022). Existing approaches to SL mainly focus on genetic interactions caused by loss-of-function events, namely by searching for lethal pairs of gene knock-outs (Thompson *et al*., 2021). This strategy has driven the development of targeted therapies, such as PARP inhibitors for patients with BRCA1- or BRCA2-mutated breast cancer (Tutt *et al*., 2021).

Despite these relevant advances, the concept of SL still requires further improvements to be more broadly applied in the growing field of precision oncology. A promising related concept is synthetic dosage lethality (SDL), which defines a genetic interaction where the overexpression of a gene combined with the under-expression of a second gene leads to cellular death (Kroll *et al*., 1996; O’Neil *et al*., 2017). In other words, SDL involves a lethal combination of one gene knock-out and one gene knock-in. The consideration of gene activation events in SDL approaches may open new therapeutic strategies, particularly with the outbreak of massively parallel knock-in CRISPR-Cas9 technologies (Roth *et al*., 2020; Dai *et al*., 2023). As in SL, SDL can be generalized to more complex interactions than 2 genes, or even to interactions exclusively involving gene knock-ins whose compound activation leads to cellular death. New and more comprehensive studies of these lethal genetic interactions are necessary to gain new insights into the identification and understanding of essential genes and tumor suppressor genes in different contexts in cancer.

Network-based computational approaches to SL have received much attention in the literature (Shen and Ideker, 2018). Specifically, given the high-quality of genome-scale metabolic networks (GEMs) of human cells (Robinson *et al*., 2020), there are a plethora of methods to predict SL and metabolic dependencies of cancel cells (Folger *et al*., 2011; Agren *et al*., 2014; Pacheco *et al*., 2019; Bintener *et al*., 2023). In previous works (Apaolaza *et al*., 2017, Apaolaza *et al*., 2019), we introduced the concept of genetic Minimal Cut Sets (gMCS), which define minimal subsets of gene knockouts that block a particular essential metabolic task, typically the biomass reaction in cancer studies. Recently, we extended our gMCS framework to go beyond cellular metabolism and integrated signed directed acyclic regulatory pathways into genome-scale metabolic networks (Barrena *et al*., 2023). We substantially increased the space of genetic interactions and uncovered new synthetic lethal candidates, leading to the hypothesis of essential genes in different cancer cell lines that were not captured exclusively with metabolic models. Importantly, this approach opens new avenues to control and target cellular proliferation in cancer cells. A full review of the gMCS approach can be found in Olaverri-Mendizabal *et al*., 2024.

In contrast with SL, SDL has not been thoroughly explored and scarce methods have been developed to identify SDLs in cancer cells (Jerby-Arnon *et al*., 2014; Megchelenbrink *et al*., 2015). A general approach was early developed by Klamt and colleagues (Klamt *et al*., 2006), who introduced the concept of Minimal Intervention Sets (MISs), which are defined as minimal set of gene knock-outs and gene knock-ins that provoke a desired response in certain molecular functions. This work was later improved by different contributions (Samaga *et al*., 2010; Garg *et al*., 2013; Biane and Delaplace, 2019; Moon *et al*., 2022). However, their application to cancer studies in large-scale molecular networks has not been further developed, due to the underlying combinatorial explosion of solutions.

Here, we present a novel methodology that extends our previous gMCS formulation to calculate genetic Minimal Intervention Sets, gMISs, which integrate interventions with both gene knockouts and knock-ins. We first show that our approach can be applied to identify gMISs in large-scale integrated metabolic and regulatory (iMR) network models. Then, using our gMIS approach, we assessed the landscape of lethal genetic interactions in human cells and the impact of gene knock-in perturbations. In addition, we applied the concept of synthetic dosage lethality to predict essential genes in cancer and evaluated its impact in sensitivity and precision with The Cancer Dependency Map (DepMap) data (Ghandi *et al*., 2019; Meyers *et al*., 2017). Moreover, we conducted a tumor suppressor analysis in cancer cell lines and identified single gene knock-in strategies that blocks cellular proliferation. Finally, we illustrate the ability of our gMIS approach to elucidate potential targets for cancer research with several examples in hematological malignancies.

## 2. Methods

In our previous work (Barrena *et al*., 2023), we presented an algorithm to calculate gMCSs in integrated networks involving metabolic and linear (acyclic) regulatory pathways, so-called here iMR networks, showing that the list of predicted SLs and genetic vulnerabilities in cancer is augmented in comparison with earlier approaches focused on the metabolic layer (Apaolaza *et al*., 2017; Valcárcel *et al*., 2024). Here, we extend this work to calculate genetic Minimal Intervention Sets, gMISs, which account not only for gene knock-out interventions, but also for interventions involving gene knock-ins.

As in our previous work (Barrena *et al*., 2023), the key innovation lies in the calculation of matrices *F* and *G*, which store minimal subsets of genetic interventions that delete a specific subset of reactions. In contrast with our previous gMCS approach, the matrices *F* and *G* involve now both gene knock-outs (*g*_*i*_^*-*^) and knock-ins interventions (*g*_*i*_^*+*^).

Figure 1 shows an example network involving 3 metabolites, 4 reactions and 7 genes, as well as the resulting matrices G and F. We obtained 6 gMISs that block the example biomass reaction (r_BIO_), among which 4 are gMCSs and the rest involves knock-in interventions, which had not been extracted with our previous gMCS approach. We present below the required modifications to calculate matrices G and F in our novel gMIS approach.

**Figure 1.**
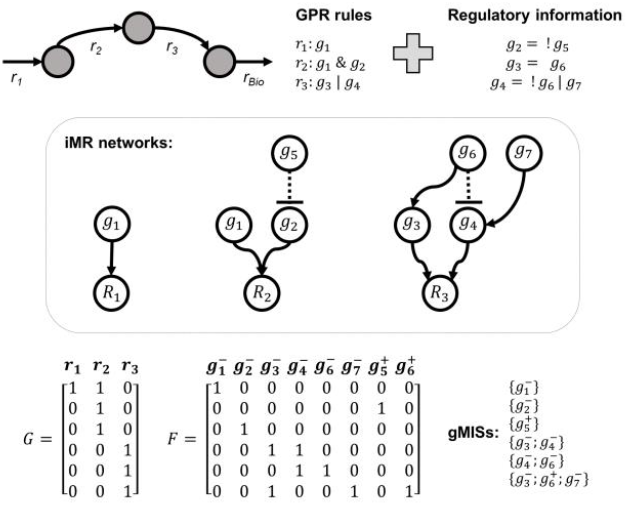
Example iMR network and gMISs. Example metabolic network and regulatory information involving 4 reactions (r_1_, r_2_, r_3_ and r_BIO_) and 7 genes (g_1_, g_2_, g_3_, g_4_, g_5_, g_6_ and g_7_). The knock-out and knock-in of gene *i* is represented as *gi*^*-*^and *gi*^*+*^, respectively.

### 2.1 Calculation of G matrix in iMR networks for gMIS calculation

As shown in Figure 1, each row in *G* defines a subset of reactions that are blocked by means of a minimal intervention set of either gene knock-outs or knock-ins, specifically detailed in its associated row in *F*. In order to build matrices *G* and *F*, we first consider each reaction separately and determine its associated minimal genetic interventions using the extended gene-protein reaction (eGPR) rules, which integrate metabolic GPR rules and regulatory pathways (see Figure 2A-B for an illustrative example). Boolean equations in eGPR rules are transformed into a linear reaction network, called eGPR network, from which we can calculate minimal genetic intervention sets using the Minimal Cut Set (MCS) approach. Finally, the set of MCSs for each reaction are integrated to build *G* and *F* matrices, as described in our previous work (Barrena *et al*., 2023). We describe below the construction of eGPR networks and the algorithm to calculate MCSs for each metabolic reaction when both gene knock-out and knock-ins are considered.

**Figure 2.**
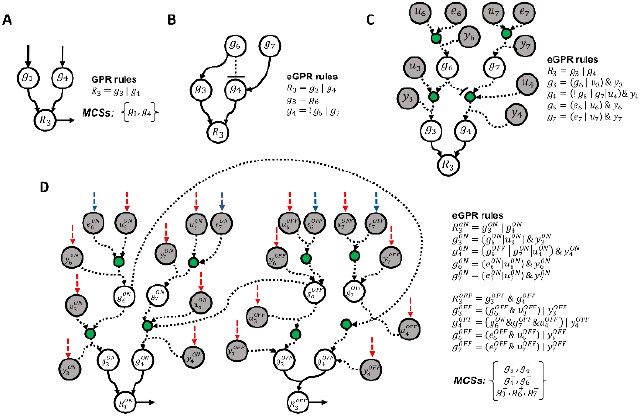
Illustration of extended GPR networks. a) GPR network for reaction 3 (*r*_*3*_) in Figure 1, its associated GPR rules and output MCSs; b) GPR rules including the regulation of metabolic genes involved in (*r*_*3*_) using Boolean equations. Two new genes are incorporated into eGPR rules: *g*_*6*_, *g*_*7*_; c) Addition of auxiliary nodes *y, u* and *e* representing gene knock-outs, gene knock-ins and external inputs, respectively, in Boolean equations; d) Resulting extended GPR (eGPR) network after dividing each node into two different ON/OFF nodes and including input and output exchange reactions. Regulatory interactions are represented through arcs in dashed line. Green nodes correspond to intermediate auxiliary nodes introduced to simplify Boolean rules. Blue arrows represent input exchange reactions for the auxiliary nodes *e* (free external inputs), while red arrows represent input exchanges associated with nodes *y* and *u* (knock-out and knock-in interventions, respectively). The network is split into ON and OFF sub-networks to systematically resolve repression logic.

#### 2.1.1 Construction of eGPR networks for gMIS calculation

For each target reaction *k*, denoted *R*_*k*_, we define *B(k)* as the subset of genes implied in its associated eGPR rule. Each of these genes, denoted *g*_*i*_ (i=1,…,|B(k)|), are interrelated by their corresponding Boolean rules. We denote *L(k)* the subset of those nodes without Boolean equations which represent input genes of the final Boolean network and can freely take 0/1 values (in Figure 2B, we have *g*_*3*_, *g*_*6*_ and *g*_*7*_). In order to build the eGPR network for each reaction, we follow 4 different steps:

i. The Boolean equation for each gene in *B(k)* is first updated with necessary auxiliary nodes *y*_*i*_ (*i*=1,…,|*B(k)*|) and *u*_*i*_ (*i*=1,…,|*B(k)*|), which allow us to consider the effect of gene knock-outs and gene knock-ins, respectively, without affecting the network upstream. Moreover, we include auxiliary nodes *e*_*i*_ (*i* ∈ *L(k)*) which allow input genes to freely take 0/1 values when no interventions are carried out. Without these nodes, we would be implicitly forcing a state on input genes even in the absence of any intervention, which would be biologically incorrect, as these nodes are meant to represent genes whose expression is free (i.e., not regulated or externally controlled in the model). The gene knock-out nodes *y*_*i*_ are added with an AND operator whereas the gene knock-in nodes *u*_*i*_ and auxiliary nodes *e*_*i*_ are added with an OR operator. The resulting Boolean network and updated eGPR rules can be found in Figure 2C. Note here that we introduce intermediate nodes (shown in green) to simplify Boolean rules.
ii. The negation terms are removed by splitting the nodes of the Boolean network from the previous step into ON and OFF nodes, namely 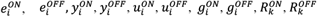 and, following the De Morgan’s laws, eGPR rules are updated. This strategy duplicates the number of nodes and interactions, but it becomes feasible to model them as a reaction network. The resulting network is shown in Figure 2D.
iii. Addition of an input exchange reaction for nodes with no input arcs, namely 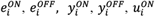 and 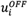 . The removal of the exchange reactions associated with 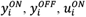 and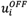, denoted here *H(k)* and colored red in Figure 2D, represents the knock-out/knock-in of the genes involved in our reaction network. In particular, the removal of the exchange reaction for 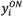 leads to the knock-out of gene *i*, whereas the removal of the exchange reaction for 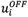 implies the knock-in of gene *i*. Clearly, they cannot occur simultaneously; however, this is restricted below in the search of MCSs described below. Finally, exchange reactions for 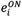 and 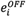 are colored blue in Figure 2D.
iv. Addition of an output exchange reaction for 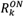 and 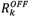, which are denoted, respectively, 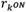 and 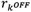 (see Figure 2D).

Nodes 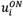 and 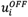 allow us to consider gene knock-ins and constitute the main difference with respect to the eGPR network built in our previous gMCS formulation.

#### 2.1.2 Calculation of MCSs in eGPR networks for gMIS calculation

eGPR networks can be modelled as a reaction system that satisfies irreversibility constraints and the mass balance equation:

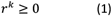

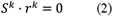

where *r*^*k*^ denotes the flux through the artificial reactions involved in the eGPR network for the target reaction *k* and *S*^*k*^ its associated stoichiometry matrix of dimensions *m*^*k*^x*n*^*k*^.

In order to calculate MCSs that block the target reaction 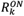, Eq. (3) forces flux through this reaction and Eq. (4) define the intervention space (gene knock-outs and knock-ins) for the input exchange reactions in *H(k):*

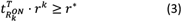

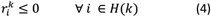

where 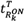 is a null vector with a 1 in the position of the target reaction 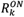 . Note here that in Eq. (4) we only include input exchange reactions in *H(k)* because they represent the decision as to whether (or not) a gene is knocked-out or knocked-in.

From the infeasible primal problem defined by Eqs. (1-4), we formulate the unbounded dual problem and minimize the number of gene knock-outs and knock-ins to block the target reaction with the following mathematical problem:

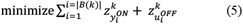

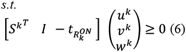

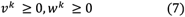

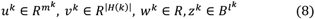

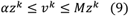

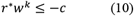

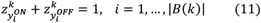

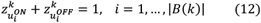

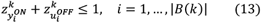

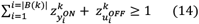

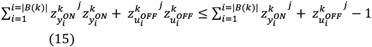

where *u, v* and *w* are dual variables associated with the mass balance equation (Eq. 2), gene intervention constrains (Eq. 4) and the target reaction (Eq. 3), respectively; *z* are the binary variables linked to *v* by Eq. (9), namely *z* = 0 ↔ *v* = 0, *z* = 1 ↔ *v* > 0. Note that here *α* and *M* are small and large positive constraints, respectively. Eq. (10) forces *w* to be non-zero, which makes the target reaction equation part of the infeasible primal problem.

The knock-out of 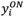 and 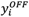 nodes, as well as 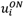 and 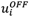 nodes, are not independent. Eq. 11 forces that for each gene *i* exactly one of these two constrains, 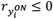 and 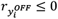, take part in the infeasible primal problem. These constraints determine that if a gene is knocked-out, *i*.*e*. 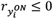, then 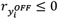 cannot be forced to be zero and vice versa. The same applies to 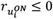 and 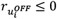 in Eq. 12, namely if a gene is knocked-in, *i*.*e*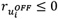, then 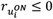 cannot be forced to be zero and vice versa. Eq. (13) establishes that we cannot knock-out and knock-in a gene *i* at the same time. Eq. (14) forces the optimal solution to involves at least one gene intervention and Eq. (15) allows us to eliminate previously obtained solutions (*z*^*j*^) from the solution space and identify new MCSs. Finally, the objective function, Eq. (5), minimizes the number of knock-outs of input exchange reactions associated with 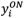 and 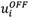, since they represent gene knock-outs 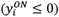 or gene knock-ins 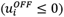. In summary, the mixed-integer linear program (MILP) defined by Eqs. (5)-allows us to enumerate MCSs for eGPR networks involving both gene knock-outs and knock-ins. With respect to our previous gMCS formulation, we have included 2 new constraints, Eqs. (12)-(13), and modified the objective function, Eq (5), and 2 constraints, Eqs. (14)-(15).

When the proposed algorithm is applied to the eGPR network of reaction 3 shown in Figure 2D, 2 solutions are obtained:

- **MCS**_**1**_: 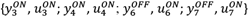
- **MCS**_**2**_: 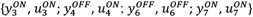

In **MCS**_**1**_, 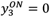indicates that g_3_ has been knocked out, and 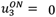 means that no knock-in has been applied to g_3_. The same applies to g_4_, where 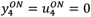. Moreover, no intervention is applied to g_6_ and g_7_, as indicated by 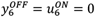 and 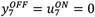 .

In **MCS**_**2**_, g_3_ and g_7_ are knocked out: 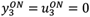and 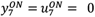. No intervention is applied to 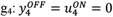 . Finally, g_6_ is knocked in: 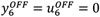.

As a result, the corresponding MCSs (in terms of gene interventions) are **MCS**_**1**_: 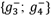 and **MCS**_**2**_: 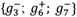. For further illustration, a more comprehensive example is provided in the Supplementary Material.

#### 2.1.3 Calculation of G matrix for gMIS calculation

We applied the mixed-integer linear program defined by Eqs. (5)-(15) to each reaction and its associated eGPR network. The resulting list of MCSs for each target reaction is then integrated to build matrices *G* and *F*, as detailed in Barrena *et al*., 2023.

### 2.2 Calculation of gMIS in iMR networks

Once the matrices *F* and *G* are adapted to consider gene knock-outs and knock-ins, the list of gMISs can be calculated using the same algorithm gMCSs presented in our previous works for gMCSs (Apaolaza *et al*., 2019; Barrena *et al*., 2023; Rodriguez-Flores *et al*., 2024). Full details can be found in Barrena *et al*., 2023.

### 2.3 Essential genes, tumor suppressors and adaptation mechanisms

With the methodology presented above, we are able to calculate gMISs in iMR networks. gMISs can include either gene knock-outs (*g*_*i*_^*-*^) or gene knock-ins (*g*_*i*_^*+*^). Following the approach presented in *gmctool* (Valcárcel *et al*., 2024), we classify a gene *g*_*i*_^*-*^as potentially essential in a particular sample if it is the unique highly expressed gene among the specific subset of gene knock-outs in a particular gMIS; however, it is also necessary to check that the subset of genes to be knocked-in in that gMIS are highly expressed. In addition, we can classify a gene *g*_*i*_^*+*^ as possibly tumor suppressor gene if it is the unique lowly expressed gene among the specific subset of gene knock-ins in a particular gMIS; however, it is also necessary to check that the subset of genes to be knocked-out in that gMIS are lowly expressed. For the definition of highly and lowly expressed genes for each sample, we applied the *gmcsTH5* threshold used in *gmctool* (Valcárcel *et al*., 2024).

Once we have identified the potential essential and tumor suppressor genes in a sample, we need to verify that when they are knocked-out or knocked-in, respectively, the rest of the genes in the gMIS of interest do not become active in the case of gene knock-outs or inactive in the case of gene knock-ins by an adaptation mechanism. To check the presence of adaptation mechanisms, we implemented a different MILP to the one presented in Barrena *et al*., 2023 (see Supplementary Material).

For illustration, Supplementary Figure 1 presents a step-by-step schematic overview of the main stages of the gMIS methodology. Finally, we validated the predicted essential genes with DepMap data. Following the same criterion presented in Robinson *et al*., 2020, a gene is deemed as essential if its essentiality score in DepMap falls below -0.6.

### 2.4 Implementation

We developed a Python function to calculate matrices *F* and *G* in iMR networks, called ‘*calculateRegNetGMatrix’*, which is available in the gMCSpy package (Rodriguez-Flores *et al*., 2024). The whole process of calculating gMISs can be done using the function ‘*calculateGMIS’* available in gMCSpy: https://github.com/PlanesLab/gMCSpy. We conducted the same implementation in MATLAB, freely available at https://github.com/PlanesLab/iMR_gMIS_paper_scripts. The code to generate all the results and the figures of the manuscript is available in 10.5281/zenodo.15350241.

For the different studies conducted in the Result section, we used the computing cluster at the University of Navarra and the Atlas supercomputer at DIPC (Donostia International Physics Center) facilities, limiting to 8 cores and 8 GB of RAM. A time limit of 5 min was set for each solution gMIS and gMCS. We made use of IBM Ilog Cplex to solve the underlying MILP model.

### 2.5 Data availability

RNA-seq and CRISPR knock-out screen data from cell lines are public and accessible in www.depmap.org (release 23Q4).

## 3. Results

In order to assess our gMIS approach, we integrated the metabolic network Human1 (v.1.14.0) (Robinson *et al*., 2020; SysBioChalmers/Human-GEM: Human 1.14.0) with the gene regulatory network of signed transcription factors DoRothEA v.1.7.2 (Garcia-Alonso *et al*., 2019), the protein-protein interaction network Omnipath (Türei *et al*., 2016) (v.3.4.7), the manually curated database of human transcriptional regulatory networks TRRUST (Han *et al*., 2018) and Signor3.0 (Surdo *et al*., 2023), a repository of manually annotated causal relationships between human proteins. The main characteristics of these regulatory and metabolic networks are shown in Supplementary Material and Supplementary Table 1.

Specifically, we built different iMR models involving 1 and 2 regulatory interaction layers for each metabolic gene: Human1 + Omnipath (Human1-O1 and Human1-O2), Human1 + DoRothEA (Human1-D1 and Human1-D2), Human1 + TRRUST (Human1-T1 and Human1-T2) and Human1 + Signor3.0 (Human1-S1 and Human1-S2). Note here that iMR models with more than 2 layers were not considered as they are strongly affected by the presence of cycles (Barrena *et al*., 2023).

For each iMR model, we calculated matrix *G* using both our gMCS and gMIS approach, this is, the *G* matrix involving only knock-out interventions (gMCS), and the *G* matrix involving both knock-out and knock-in interventions (gMIS). Following the recent work of Olaverri-Mendizabal *et al*., 2024 (Olaverri-Mendizabal *et al*., 2024), where it is shown that the contribution of gMCSs larger than 5 genes to the prediction of genetic vulnerabilities is negligible, we decided to simplify matrix *G* and discard rows involving more than 5 genes. In addition, we computed the set of gMCSs and gMISs until length 5 that block the biomass production for each iMR model. Full details of the computational performance are provided in Supplementary Table 2, showing that the computation time scales linearly with the number of rows of the G matrix (Pearson′s correlation = 0.98, *p value* = 2.1e-11), which demonstrates the efficiency and the scalability of gMIS in comparison the original gMCS algorithm. Supplementary Figures 2-3 summarize the landscape of lethal genetic interactions for the different regulatory databases considered and show the clear increment of gMISs with respect to gMCSs.

We validated the predicted gMCSs and gMISs using the SLKB database (Gökbağ *et al*., 2024), which includes 16,059 experimentally validated SL gene pairs and 264,424 non-SL pairs from 11 combinatorial knockout studies across 22 cancer cell lines. Our results showed statistically significant enrichment between gMCS/gMIS predictions and SLKB entries in all iMR models tested. Notably, the highest p-value observed among all models was 0.00225 (Human1-O2-SDL), with the rest of models yielding even smaller p-values. These findings highlight the robustness and biological relevance of our approach.

### 3.1 Prediction of essential genes in cancer based on SL and SDL interactions

By combining the calculated gMISs and gene expression from RNA-seq data, we can predict context-specific essential genes in cancer, as described in Methods section. Using gene expression data from CCLE (Ghandi *et al*., 2019), we computed a list of essential genes per cell line and iMR model in two different scenarios: i) considering only SL interactions (gMCSs); and ii) considering both SL and SDL interactions (gMCSs and gMISs). In the first case, a target gene is predicted as essential if it takes part of at least one gMCS where the rest of the genes to be knocked-out are lowly expressed; in the second case, a target gene is predicted as essential if it takes part of at least one gMIS where the rest of the genes to be knocked-out are lowly expressed and the subset of genes to be knocked-in are highly expressed. We checked in both cases that lowly expressed genes do not become highly expressed and vice versa upon the knock-out of the target gene (see Supplementary Material), due to a potential adaptation mechanism according to their associated Boolean regulatory rules. Considering the set of 1,021 cancer cell lines from CCLE, Figure 3A shows the impact of SDL interactions in the number of predicted essential genes when compared to SL interactions that are not present in Human1. It can be observed that the number of essential genes is increased in all cases, particularly in Human1-O1 (one-sided paired Wilcoxon test p-value < 2.2e-16, mean difference of essential genes per cell line = 8.007), Human1-O2 (one-sided paired Wilcoxon test p-value < 2.2e-16, mean difference of essential genes per cell line = 7.211) and Human1-T2 (one-sided Wilcoxon test p-value < 2.2e-16, mean difference of essential genes per cell line = 3.917), which illustrates the relevance of SDL interactions in the prediction of genetic vulnerabilities. Of note, the iMR models built on Signor3.0 are the ones where SDL interactions have less influence in the number of essential genes (mean difference of essential genes per cell line ≤ 0.582). The smaller effect of SDL interactions in Signor3.0-based iMR models can be explained by the lower number of gMIS involving gene knock-ins. For instance, while Human1-S2 presents 143 gMISs involving gene knock-ins, Human1-O2 and Human1-T2 show 1,397 and 11,250 such cases, respectively. These differences also reflect the distinct nature of the biological interactions of the underlying databases: Signor3.0 encodes protein–protein interaction relationships, whereas TRRUST and DoRothEA capture transcription factor–target relationships.

**Figure 3.**
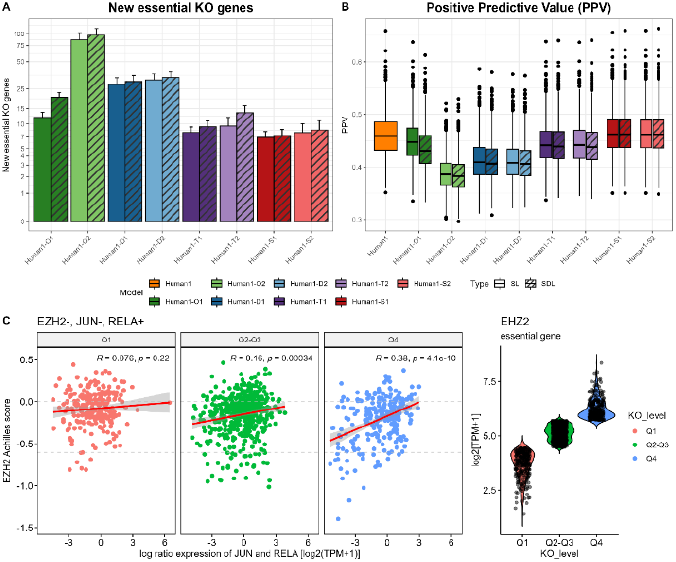
Gene essentiality analysis of iMR models with the gMCS and gMIS approaches. A) Average number of essential genes across cancer cell lines obtained from gMCSs and gMISs of different iMR models; B) Boxplot of Positive Predictive Value (PPV) of different cancer cell lines using DepMap data as a gold-standard of gene essentiality obtained from the gMCS and gMIS approaches; C) Correlation between the essentiality of EZH2 (CRISPR knock-out screen data from DepMap) and the difference in expression between JUN and RELA in log2(TPM+1) in all DepMap cell lines in three cases: i) cell lines where the expression of EZH2 falls within the first quartile (Q1), ii) the second and third quartile (Q2-Q3) and iii) the fourth quartile (Q4). The gMIS approach predicts that EZH2 is more essential when the JUN/RELA expression ratio is low, i.e. when RELA expression is high and JUN expression is low. This pattern is more evident under high EZH2 expression (Q4), where a stronger correlation between EZH2 essentiality and the JUN/RELA ratio is observed, compared to conditions of low (Q1) or moderate (Q2–Q3) EZH2 expression.

In addition, we compared in both cases the predicted essential genes with DepMap essentiality scores (Meyers et al., 2017) (see Methods section). In particular, for each iMR model, we estimated the positive predicted value (PPV) (Figure 3B), which is defined as the ratio of true positives to all the genes defined as positives (true and false positives) (see Supplementary Table 3 for details). The most accurate results were found in Human1-S1 and Human-S2, slightly superior to those of Human1. In particular, the highest PPV value was found in Human1-S1 and the gMIS approach (PPV=0.465). In addition, aside from Human1-O1, where the inclusion of SDL interactions worsens the prediction of essential genes (one-sided paired Wilcoxon test p-value < 2.2e-16, mean difference of PPV = 1.4%), the resulting PPV in the rest of the cases remains very similar (one-sided paired Wilcoxon test p-value < 2.2e-16, mean difference of PPV per cell line ≤ 0.17%). For completeness, we also analyzed false negatives and model sensitivity (SE) for different iMR models (Supplementary Table 3). Consistently, the gMIS approach showed a slight but consistent increase in SE relative to the gMCS approach (one-sided paired Wilcoxon test p-value < 2.2e-16, mean difference of SE per cell line ≥ 0.18%).

Finally, we analyzed the list of essential genes and SDL interactions predicted with our gMIS approach in the different iMR models (full details in Supplementary Table 4). Among different candidates, we identified the essentiality of EZH2 specifically in hematological cancer cell lines (Supplementary Figure 4) in all iMR models build on TRRUST. Interestingly, this gene has been previously identified as a drug target of different hematological tumors (Herviou *et al*., 2016). Here, the lethality of EZH2 inhibition is mediated by the following gMIS: {EZH2^-^, JUN^-^, RELA^+^}, hypothesizing a higher impact in cellular proliferation when the expression of JUN is down-regulated and RELA is up-regulated. We found a good association with DepMap data (Fisher’s test *p-value*: 1.192e-05, odds ratio: 3.705), particularly in lymphoid cell lines (Supplementary Figure 4). In addition, we found a significant positive correlation between the DepMap essentiality scores of EZH2 and the difference in gene expression of JUN and RELA in the set of cell lines analyzed, in line with our theoretical prediction (Supplementary Figure 5). However, we observed that this positive correlation is strongly dependent on the expression of EZH2 (Figure 3C). Specifically, little correlation was found for cell lines where the expression of EZH2 falls within the first quartile, Q1, (Pearson Correlation test p-value=0.22, r=0.076), but a higher correlation for those cell lines where the expression of EZH2 is among the second and third quartile, Q2-Q3, (Pearson Correlation test p-value = 0.00034, r=0.16) and the fourth quartile, Q4, (Pearson Correlation test p-value = 4.1e-10, r=0.35). This indicates that the effect of EZH2 inhibition is more clearly observed in cell lines where it is highly expressed.

### 3.2 Tumor suppressors analysis with the gMIS approach

In contrast with our previous gMCS approach, the gMIS approach allows us to predict single gene knock-in strategies that blocks cellular proliferation. They can be directly obtained from global tumor suppressor genes (gMISs of length 1 with 1 gene knock-in), but also from gMISs of length > 1, particularly from SDL interactions and tumor suppressor gene complexes (gMIS of length 1 only comprising knock-in interventions), leading to context-specific interventions. We describe below how context-specific lethal gene knock-ins are inferred with our gMIS approach.

For illustration, consider the SDL interaction formed by the following 2 genes: {DUSP4^+^; GADD45B^-^}, where the high expression of DUSP4 and the low expression of GADD45B inhibit cellular proliferation. Put differently, the low expression values of GADD45B indicates that the knock-in of DUSP4 is lethal for cellular proliferation. Thus, DUSP4 is a context-specific tumor suppressor that is dependent on the activity of GADD45B. In the absence of large-scale gene knock-in screenings, we made use of DepMap data to show the opposite situation: higher cellular proliferation upon the gene knock-out of DUSP4 is found for higher expression values of GADD45B (Supplementary Figure 6), being particularly true for cell lines where DUSP4 is highly expressed (Q4 in Figure 4A, Pearson Correlation test p-value = 2.5e-6, r=0.29).

For completeness, we also examined the following tumor suppressor gene complex comprising 2 genes: {CDK5RAP3^+^; SHC1^+^}, where the high expression of both genes is required to block cellular proliferation and, thus, if one of them is lowly expressed, its knock-in is lethal for cellular proliferation. As done above, we confirmed the opposite situation based on DepMap data: higher cellular proliferation upon the gene knock-out of CDK5RAP3 is found for lower expression values of SHC1 (Supplementary Figure 7). Again, this is more clearly observed for cell lines where CDK5RAP3 is highly expressed (Q4 in Figure 4B, Pearson Correlation test p-value = 3.1e-5, r=-0.26).

**Figure 4.**
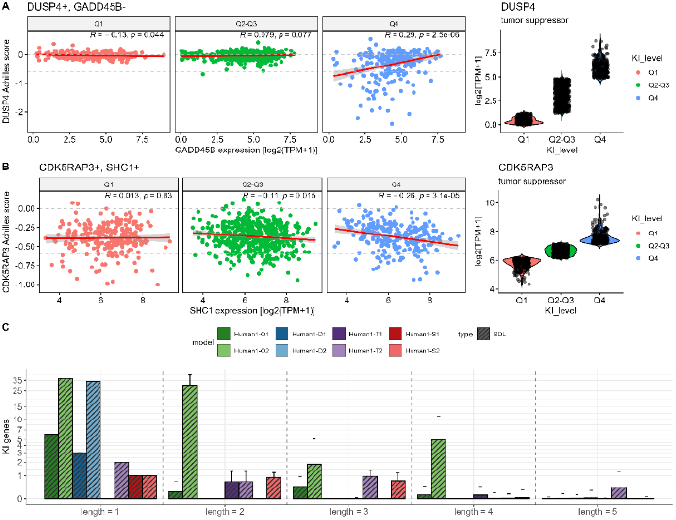
Tumor suppressor gene analysis for different iMR models with the gMIS approach. A) Correlation between the essentiality of DUSP4 (CRISPR knock-out screen data from DepMap) and the expression of GADD45B in log2(TPM+1) in all DepMap cell lines in three cases: i) cell lines where the expression of DUSP4 falls within the first quartile (Q1), ii) the second and third quartile (Q2-Q3) and iii) the fourth quartile (Q4). The gMIS approach predicts that DUSP4 is less essential when GADD45B expression is high. This is more clearly observed when DUSP4 expression is high (Q4), as indicated by a stronger positive correlation between DUSP4 essentiality and the GADD45B expression compared to conditions of low (Q1) or moderate (Q2–Q3) DUSP4 expression; B) Correlation between the essentiality of CDK5RAP3 (CRISPR knock-out screen data from DepMap) and the expression of SHC1 in log2(TPM+1) in all DepMap cell lines in three cases: i) cell lines where the expression of CDK5RAP3 falls within the first quartile (Q1), ii) the second and third quartile (Q2-Q3) and iii) the fourth quartile (Q4). The gMIS approach predicts that CDK5RAP3 is less essential when the SHC1 expression is low. The predictive logic of the gMIS approach becomes more apparent when CDK5RAP3 expression is high (Q4), as indicated by a stronger negative correlation between CDK5RAP3 essentiality and the SHC1 expression compared to conditions of low (Q1) or moderate (Q2–Q3) DUSP4 expression; C) Average number of tumor suppressor genes across cancer cell lines obtained from gMISs of different lengths (1,2,3,4,5) in different iMR models.

Using this computational framework, we predicted lethal gene knock-in strategies for different iMR models (Figure 4C). We found the highest number of interventions in the models built on Omnipath, finding an average number of tumor suppressor genes per cell line in Human1-O1 and Human1-O2 of 6.79 and 68.54, respectively. At the other extreme, we have Human1-S1 and Human1-S2, which only obtained an average number of solutions of 1.016 and 1.945, respectively.

In order to validate our tumor suppressor gene predictions, we used the TSGene2.0 database (Zhao *et al*., 2016), which has information for 1217 tumor suppressor genes curated from more than 9000 articles. We found a very significant enrichment of our tumor suppressor predictions in Human-O2 (one-sided hypergeometric test p-value = 3.02e-11), Human-D1 (one-sided hypergeometric test p-value = 1.5e-2), Human-D2 (one-sided hypergeometric test p-value = 1.05e-11), Human-T1 (one-sided hypergeometric test p-value = 2.7e-3) and Human-T2 (one-sided hypergeometric test p-value = 1.86e-06) with TSGene database, validating the predictive capability of our approach (Supplementary Table 5 for details).

## 4. Discussion

In this work, we present a novel computational method that predicts lethal genetic interactions within integrated metabolic and regulatory (iMR) networks of human cells. Specifically, we expand on our previous genetic Minimal Cut Set (gMCS) formulation to compute genetic Minimal Intervention Sets (gMISs), which incorporate both gene knock-out and knock-in perturbations. We show that our approach is capable of analyzing large-scale iMR network models and identifying different type of lethal genetic interactions: global essential genes, synthetic lethality, global tumor suppressor genes, synthetic dosage lethality (SDL) and tumor suppressor gene complexes (TSGC). To the best of our knowledge, this is the first constraint-based modeling approach able to systematically uncover these interactions in human cells, with a particular focus on SDL, which has received limited attention in the literature, and TSGC, a concept that is newly introduced in this work. Note here that previous methods, such as Mahadevan *et al*., 2015 and Schneider *et al*., 2022, have incorporated interventions that up- or down-regulate reactions rates. Our approach is fundamentally different, extending beyond cellular metabolism by integrating linear regulatory and signaling pathways, which involve inhibition rules.

Our gMIS approach was successfully implemented on 4 different iMR networks with 1 and 2 regulatory layers: Human1 with Omnipath (Human1-O1, Human1-O2), DoRothEA (Human1-D1, Human1-D2), TRRUST (Human1-T1, Human1-T2) or Signor3.0 (Human1-S1, Human1-S2). We computed solutions up to length 5 for all these iMR models and observed significant variability across regulatory databases. For instance, Human1-T2 yielded 22,210 gMISs and Human1-D2 11,449 gMISs (Supplementary Table 2). This highlights the importance of creating consensus models, similar to what is typically done in genome-scale metabolic models.

As in our previous gMCS approach, RNA-seq data can be mapped onto our database of gMISs to predict cancer-specific single gene knock-outs that are lethal for cellular proliferation. Here, using large-scale single gene knock-out screens from DepMap, we show that SDL allows us to extract context-specific essential genes that were not recovered solely with synthetic lethality interactions. We showcase our methodology with EZH2, a component of the polycomb-group protein complexes (Liu and Yang, 2023), which functions by methylating histones on chromatin to inhibit the transcription initiation of target genes. Consistent with DepMap findings, EZH2 is found to play a crucial role in hematological malignancies. In contrast with our previous gMCS approach, RNA-seq data can be integrated with our database of gMISs to predict cancer-specific single gene knock-ins that are lethal for cellular proliferation, *i*.*e*. cancer-specific tumor suppressor genes. The primary challenge in validating our computational predictions lies in the scarcity of large-scale gene knock-in screens in the existing literature. Although new CRISPR-Cas9 knock-in technologies are emerging (Roth *et al*., 2020; Dai *et al*., 2023), they are still in their infancy compared to gene knock-out screens like DepMap, which are very well established. We partially circumvented this issue by using DepMap data under the assumption that the knockout of a tumor suppressor gene results in an increase in cellular proliferation, leading to a higher Achilles score, as observed in Figure 4A-B with 2 different examples. We acknowledge that this validation is incomplete, and the availability of large-scale gene knock-in screen data is essential to conclusively validate our predictions. However, the results obtained with TSGene database is promising.

Finally, even though our gMIS approach currently cannot accommodate cyclic Boolean rules, which restricts the integration of metabolic and regulatory layers, the proposed methodology paves the way for mechanistically predicting novel genetic interactions and metabolic dependencies in tumor cells. The computational and functional biological analysis presented here demonstrates that gMIS approach can be effectively utilized to investigate the regulation of cancer metabolism and its related dependencies.

## Acknowledgements

This work was supported by the Minister of Economy and Competitiveness of Spain [PID2022-143298OB-I00, F.J.P.] and PIBA Programme of the Basque Government [PIBA_2025_1_0048, F.J.P.]. N.B. received his salary from Basque Government predoctoral grant [PRE_2023_2_0158]. The funders had no role in study design, data collection and analysis, decision to publish, or preparation of the manuscript.

## Supplementary data

Supplementary data are available at *Bioinformatics Advances* online.

