## Supplementary Material for "Beyond synthetic lethality in large-scale metabolic and regulatory network models via genetic minimal intervention sets"

### Regulatory and metabolic networks of human cells

To assess our novel approach, we employed the protein-protein interaction network of Omnipath (Türei *et al.*, 2016) (accessed online 2023-04-03) (OmnipathR, v.3.0.4), the gene regulatory network of signed transcription factors DoRothEA (Garcia-Alonso *et al.*, 2019) (dorothea, v.1.7.2), the manually curated database of human transcriptional regulatory networks TRRUST (Han *et al.*, 2018) and Signor3.0 (Surdo *et al.*, 2023) (Signor, v.3.0). Specifically, for the database Signor3.0, we only selected interactions with evidence score ≥ 0.5. The main characteristics of each regulatory network are shown in Table 1.

Regarding to the metabolic model, we used the most recent genome-scale metabolic network of human cells at the time of the onset of study: Human1 (Robinson *et al.*, 2020; SysBioChalmers/Human-GEM: Human 1.14.0) (v1.14.0) obtained from https://github.com/SysBioChalmers/Human-GEM. Human1 involves 8,363 metabolites, 2,920 genes and 13,024 reactions. Although this model defines 56 essential metabolic tasks, for simplicity, we focus here on the task of producing biomass under the Ham’s medium.

Human1 makes use of Ham’s medium to produce biomass. Therefore, the flux through the input exchange reactions of metabolites not involved in Ham’s medium, as defined in Human1 for biomass production, was set to zero. Then, the model was simplified with the function *simplifyModel* of RAVEN (Wang *et al.*, 2018), deleting reactions that are constrained to zero flux. After this simplification, Human1 is reduced to 6,830 metabolites, 2,419 genes and 11,573 reactions.

In order to generate integrated metabolic and regulatory networks, we add signed interactions for each metabolic gene involved in each GPR of the metabolic network. Then, we create a new Boolean equation that integrates the interactions for each metabolic gene using ‘OR’ operators building eGPR rules.

As in Barrena *et al*., 2023 (Barrena *et al.*, 2023), our gMIS approach is not able to deal with cyclic behaviors in Boolean networks. To avoid cycles in eGPR rules, at the time of adding a regulatory layer, we followed the same procedure presented in Barrena *et al*., 2023 (Barrena *et al.*, 2023), where the presence of cycles is checked and regulatory interactions are only included if the assessed eGPR rules is cycle free.

**Adaptation mechanisms in the prediction of gene essentiality**

Once we have identified the potential essential and tumor suppressor genes in a sample, we need to verify that when they are knocked-out or knocked-in, respectively, the rest of the genes in the gMIS of interest do not become active in the case of gene knock-outs or inactive in the case of gene knock-ins by an adaptation mechanism. To check the presence of adaptation mechanisms, we implement a similar MILP to the one presented in Barrena et al., 2023.

To check the presence of adaptation mechanisms, we independently transform the Boolean equations of each eGPR into linear constraints with binary variables *x*, similarly to the work presented in Shlomi et al., 2007 (Shlomi *et al.*, 2007). Note here that we only need to consider eGPRs that contain at least one gene of the target gMIS. We add auxiliary nodes *y* and *u* in eGPR rules to model gene knock-outs and knock-ins, respectively, as explained above in the construction of the eGPRs in the main text. Then, we force the knock-out of the potential essential gene T ($x_{Y_{T}}=0$) or the knock-in of the potential tumor suppressor T ($x_{U_{T}}=1$) and minimize the number of gene expression changes for each eGRP independently. Specifically, we give a negative coefficient in the objective function to gene knock-ins ($x_{j}^{+}$) to promote their activation. The opposite applies to gene knock-outs ($x_{j}^{-}$). We solve this problem by integer linear programming:

$\mathrm{minimize}\sum_{j\in P_{T}} x_{j}^{-}-x_{j}^{+}$	(16)

*s.t.*

$\alpha\leq A\cdot x\leq\beta$	(17)

$x_{Y_{T}}=0 | x_{U_{T}}=1$	(18)

$x:\left\{ 0,1 \right\}$	(19)

If the objective value of the above integer linear programming is equal to the number of genes to be knocked-in the gMIS with negative sign, tentative essential genes or tumor suppressors are kept. If the objective function takes a greater value than the above, an adaptation mechanism is assumed and tentative essential genes or tumor suppressors are discarded.

**Constructed eGPR and Computed MCSs for the Reaction MAR04379 in the Model Human1-T1-SDL**

To illustrate the application of the MILP formulation (Eqs. (5)-(15) in the main manuscript), we present a case study based on reaction MAR04379 from the Human1 model.

Step 1: eGPR Construction

The original metabolic GPR for MAR04379 is ‘ENSG00000067057 or ENSG00000141959 or ENSG00000152556’, which corresponds to PFKP OR PFKL OR PFKM.

Using transcriptional regulation data from the TRRUST database, the following regulatory rule is added:

- $PFKP = KLF4 | !ZBTB7A$

The resulting extended GPR (eGPR) rules are:

- $R_{MAR04379}= PFKP | PFKL | PFKM$
- $PFKP = KLF4 | !ZBTB7A$

Step 2: Addition of Auxiliary Nodes

Auxiliary variables are added to encode gene knock-ins (u), knock-outs (y), and auxiliary input nodes (e):

- $R_{MAR04379}= PFKP | PFKL | PFKM$
- $PFKP = (KLF4 | !ZBTB7A | u_{PFKP}) \& y_{PFKP}$
- $PFKL = \left( e_{PFKL} \right|u_{PFKL}) \& y_{PFKL}$
- $PFKM = \left( e_{PFKM} \right|u_{PFKM}) \& y_{PFKM}$
- $KLF4 = \left( e_{KLF4} \right|u_{KLF4}) \& y_{KLF4}$
- $ZBTB7A = \left( e_{ZBTB7A} \right|u_{ZBTB7A}) \& y_{ZBTB7A}$

Step 3: ON/OFF Node Decomposition

Negations are eliminated by decomposing all nodes into ON and OFF states.

ON configurations:

- $R_{MAR04379}^{ON}= PFKP^{ON} \left| PFKL^{ON} \right|PFKM^{ON}$
- $PFKP^{ON} = \left( KLF4^{ON} \right|ZBTB7A^{OFF} | u_{PFKP}^{ON}) \& y_{PFKP}^{ON}$
- $PFKL^{ON} = \left( e_{PFKL}^{ON} \right|u_{PFKL}^{ON}) \& y_{PFKL}^{ON}$
- $PFKM^{ON} = \left( e_{PFKM}^{ON} \right|u_{PFKM}^{ON}) \& y_{PFKM}^{ON}$
- $KLF4^{ON} = \left( e_{KLF4}^{ON} \right|u_{KLF4}^{ON}) \& y_{KLF4}^{ON}$
- $ZBTB7A^{ON} = \left( e_{ZBTB7A}^{ON} \right|u_{ZBTB7A}^{ON}) \& y_{ZBTB7A}^{ON}$

OFF configurations:

- $R_{MAR04379}^{OFF}= PFKP^{OFF} \& PFKL^{OFF}\& PFKM^{OFF}$
- $PFKP^{OFF} =\left( KLF4^{OFF} \& ZBTB7A^{ON} \& u_{PFKP}^{OFF} \right) | y_{PFKP}^{OFF}$
- $PFKL^{OFF} =\left( e_{PFKL}^{OFF} \& u_{PFKL}^{OFF} \right) | y_{PFKL}^{OFF}$
- $PFKM^{OFF} =\left( e_{PFKM}^{OFF}\& u_{PFKM}^{OFF} \right) | y_{PFKM}^{OFF}$
- $KLF4^{OFF} =\left( e_{KLF4}^{OFF}\& u_{KLF4}^{OFF} \right) | y_{KLF4}^{OFF}$
- $ZBTB7A^{OFF} =\left( e_{ZBTB7A}^{OFF}\& u_{ZBTB7A}^{OFF} \right) | y_{ZBTB7A}^{OFF}$

Step 4: MILP Formulation

To compute the MCSs for this eGPR network, we solve the following MILP (from Eqs. (5)-(15) in the manuscript):

$\mathrm{minimize}\sum_{i=1}^{i=|B(k)|} z_{y_{i}^{ON}}^{k}+ z_{u_{i}^{OFF}}^{k}$	(1)

*s.t.*

$\left[ {S^{k}}^{T} I -t_{R_{k}^{ON}} \right] \left( \begin{matrix} u^{k} \\ v^{k} \\ w^{k} \end{matrix} \right)\geq0$	(2)

$v^{k} \geq0, w^{k} \geq0$	(3)

$u^{k}\in R^{m^{k}}, v^{k}\in R^{\left| H\left( k \right) \right|}, w^{k}\in R, z^{k}\in B^{l^{k}}$	(4)

$\alpha z^{k}\leq v^{k}\leq Mz^{k}$	(5)

$r^{*}w^{k}\leq-c$	(6)

$z_{y_{i}^{ON}}^{k}+z_{y_{i}^{OFF}}^{k}=1, i=1,\ldots,\left| B\left( k \right) \right|$	(7)

$z_{u_{i}^{ON}}^{k}+z_{u_{i}^{OFF}}^{k}=1, i=1,\ldots,\left| B\left( k \right) \right|$	(8)

$z_{y_{i}^{ON}}^{k}+z_{u_{i}^{OFF}}^{k}\leq1, i=1,\ldots,\left| B\left( k \right) \right|$	(9)

$\sum_{i=1}^{i=|B(k)|} z_{y_{i}^{ON}}^{k}+ z_{u_{i}^{OFF}}^{k}\geq1$	(10)

$\sum_{i=1}^{i=|B(k)|} {z_{y_{i}^{ON}}^{k}}^{j}z_{y_{i}^{ON}}^{k}+ {z_{u_{i}^{OFF}}^{k}}^{j}z_{u_{i}^{OFF}}^{k}\leq\sum_{i=1}^{i=|B(k)|} {z_{y_{i}^{ON}}^{k}}^{j}+ {z_{u_{i}^{OFF}}^{k}}^{j}-1$	(11)

- Eqs. (1)-(6): Define the dual of the FBA problem and optimize to find minimal knock-out/knock-in interventions.
- Eq. (7)-(8): Enforce mutual exclusivity between ON/OFF states of each node.
- Eq. (9): Avoid simultaneous knock-out and knock-in for a given gene.
- Eq. (10): Ensure that at least one gMIS is found per MILP run.
- Eq. (11): Exclude previously found solutions to allow for enumeration of new MCSs.

For example, Eqs. (7)-(9) applied to the node PFKP reads:

$$z_{y_{PFKP}^{ON}}+z_{y_{PFKP}^{OFF}}=1$$

$$z_{u_{PFKP}^{ON}}+z_{u_{PFKP}^{OFF}}=1$$

$$z_{y_{PFKP}^{ON}}+z_{u_{PFKP}^{OFF}}\leq1$$

and Eq. 10:

$$z_{y_{PFKP}^{ON}}+z_{u_{PFKP}^{OFF}}+ z_{y_{PFKL}^{ON}}+z_{u_{PFKL}^{OFF}}+ z_{y_{PFKM}^{ON}}+z_{u_{PFKM}^{OFF}}+ z_{y_{KLF4}^{ON}}+z_{u_{KLF4}^{OFF}}+ z_{y_{ZBTB7A}^{ON}}+z_{u_{ZBTB7A}^{OFF}}\geq1$$

Step 5: Resulting MCSs

After solving the MILP for this eGPR configuration, we identify the following gMCSs:

**MCS1** (gene-level):

{$y_{PFKP}^{ON}$,$u_{PFKP}^{ON}$;$y_{PFKL}^{ON}$,$u_{PFKL}^{ON}y_{PFKM}^{ON}$,$u_{PFKM}^{ON}$; $y_{KLF4}^{OFF}$,$u_{KLF4}^{ON}$;$y_{ZBTB7A}^{OFF}$,$u_{ZBTB7A}^{ON}$}

**MCS2** (gene-level):

{$y_{PFKP}^{OFF}$,$u_{PFKP}^{ON}$;$y_{PFKL}^{ON}$,$u_{PFKL}^{ON}y_{PFKM}^{ON}$,$u_{PFKM}^{ON}$; $y_{KLF4}^{ON}$,$u_{KLF4}^{ON}$;$y_{ZBTB7A}^{OFF}$,$u_{ZBTB7A}^{OFF}$}

These correspond to the following **gene-level interventions**:

- **MCS_1_**: {$g_{PFKP}^{-}$; $g_{PFKL}^{-}$; $g_{PFKM}^{-}$}
- **MCS_2_**: {$g_{PFKL}^{-}$; $g_{PFKM}^{-}$; $g_{KLF4}^{-}$; $g_{ZBTB7A}^{+}$}

**Supplementary Figures**

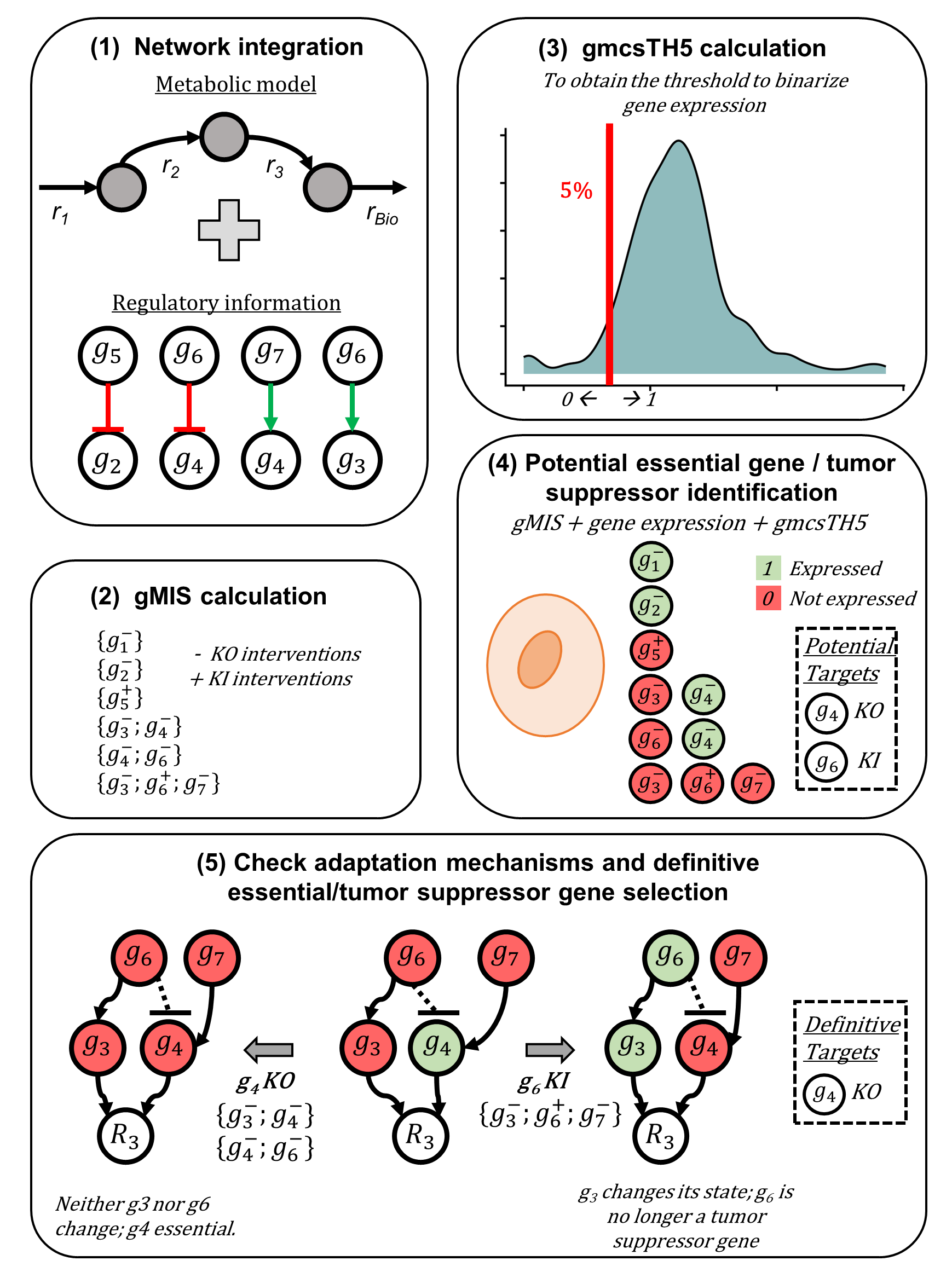

**Supplementary Figure 1.** **Overview of the gMIS workflow**. Schematic representation of the main steps of the gMIS methodology. (1) **Network integration:** metabolic and regulatory networks are combined to generate an integrated model. (2) **gMIS calculation:** genetic Minimal Intervention Sets (gMISs) are computed. gMISs include both gene knock-out (KO) and knock-in (KI) interventions. (3) **gmcsTH5 calculation: gmcsTH5 is computed to** binarize gene expression and integrate transcriptomic data with the gMIS (Valcárcel *et al.*, 2024). (4) **Potential essential gene and tumor suppressor identification:** potential essential and tumor suppressor genes are predicted by combining the gMIS with gene expression data and the gmcsTH5 threshold, distinguishing expressed (green) and non-expressed (red) genes. (5) **Adaptation mechanisms check and definitive target selection:** Adaptation regulatory mechanisms are evaluated to confirm final essential or tumor suppressor genes.

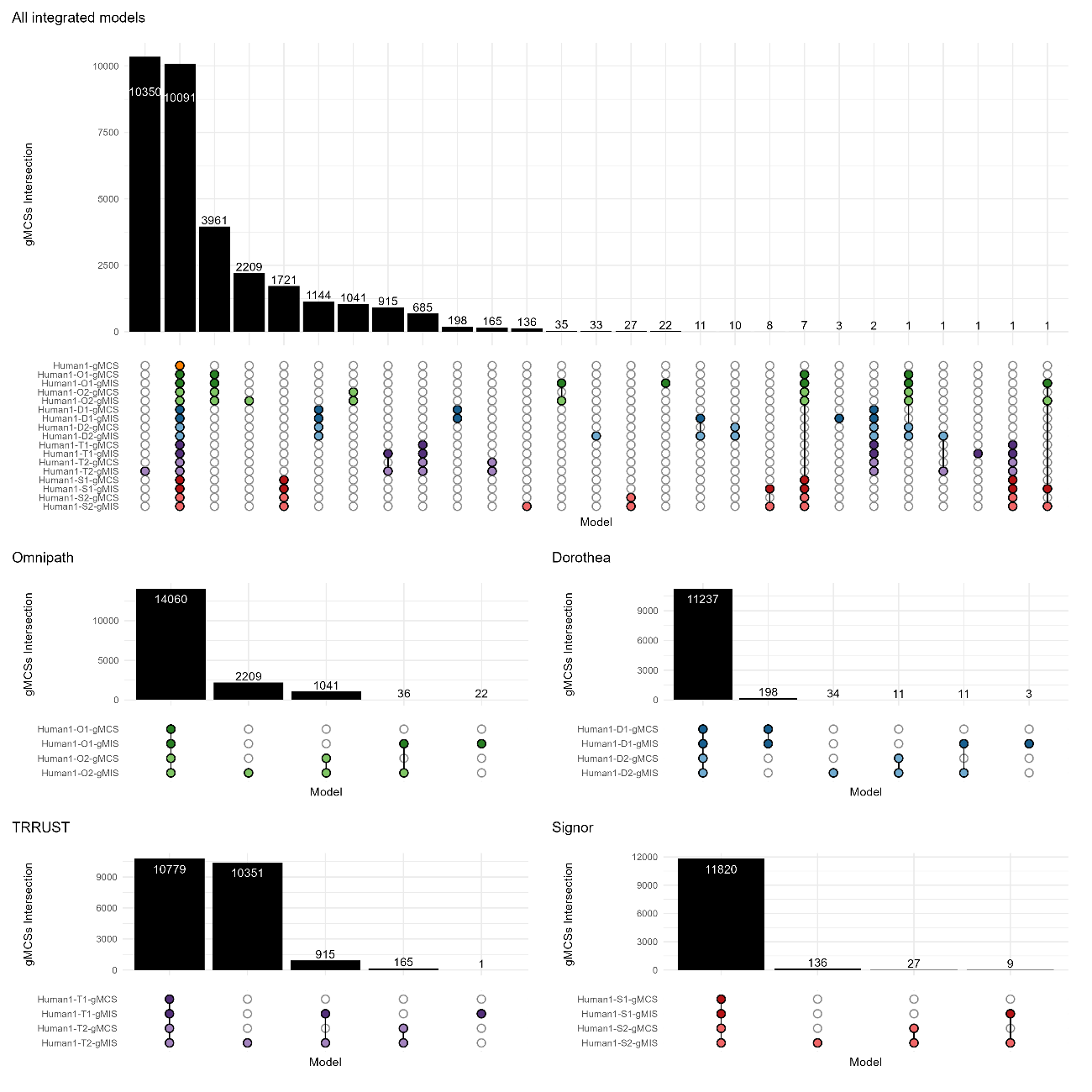

**Supplementary Figure 2.** **Analysis of gMCS and gMIS obtained from different integrated metabolic and regulatory networks**. Upset plots representing the intersection of the gMCSs and gMISs obtained from the different integrated metabolic and regulatory networks: Human1, Human1-O1, Human1-O2, Human1-D1, Human1-D2, Human1-T1, Human1-T2, Human1-S1, Human1-S2. Abbreviations: ‘*Human1-O1’*: integrated model with Human1 and Omnipath with one regulatory layer; ‘*Human1-O2’*: integrated model with Human1 and Omnipath with two regulatory layers; ‘*Human1-D1’*: integrated model with Human1 and DoRothEA with one regulatory layer; ‘*Human1-D2’*: integrated model with Human1 and DoRothEA with two regulatory layers; ‘*Human1-T1’*: integrated model with Human1 and TRRUST with one regulatory layer; ; ‘*Human1-T2’*: integrated model with Human1 and TRRUST with two regulatory layers; ‘*Human1-S1’*: integrated model with Human1 and Signor3.0 with one regulatory layer; ‘*Human1-S2’*: integrated model with Human1 and Signor3.0 with two regulatory layers.

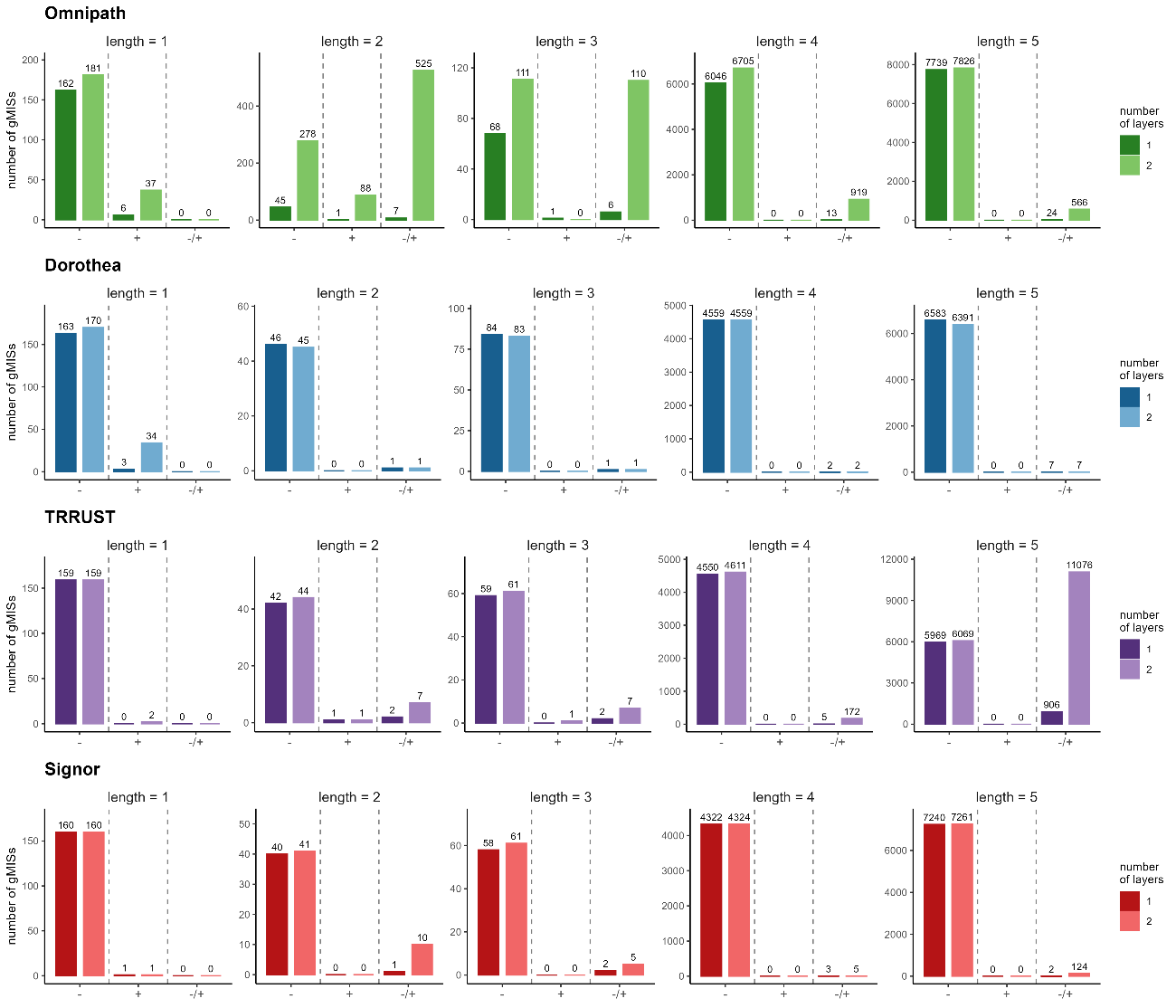

**Supplementary Figure 3. Number and length of gMISs for different iMR models.** The negative symbol ‘-’ refers to gMISs involving only gene knock-out interventions, *i.e.* gMCSs. Solutions of length 1 are global essential genes, and solutions of length > 1 are synthetic lethals. The positive symbol ‘+’ refers to gMISs involving only gene knock-in interventions. Solutions of length 1 are global tumor suppressor genes, and solutions with length > 1 are called tumor suppressor gene complexes; and the symbol ‘-/+’ refers to gMISs involving both gene knock-out and knock-in interventions, which define synthetic dosage lethal interactions of different length.

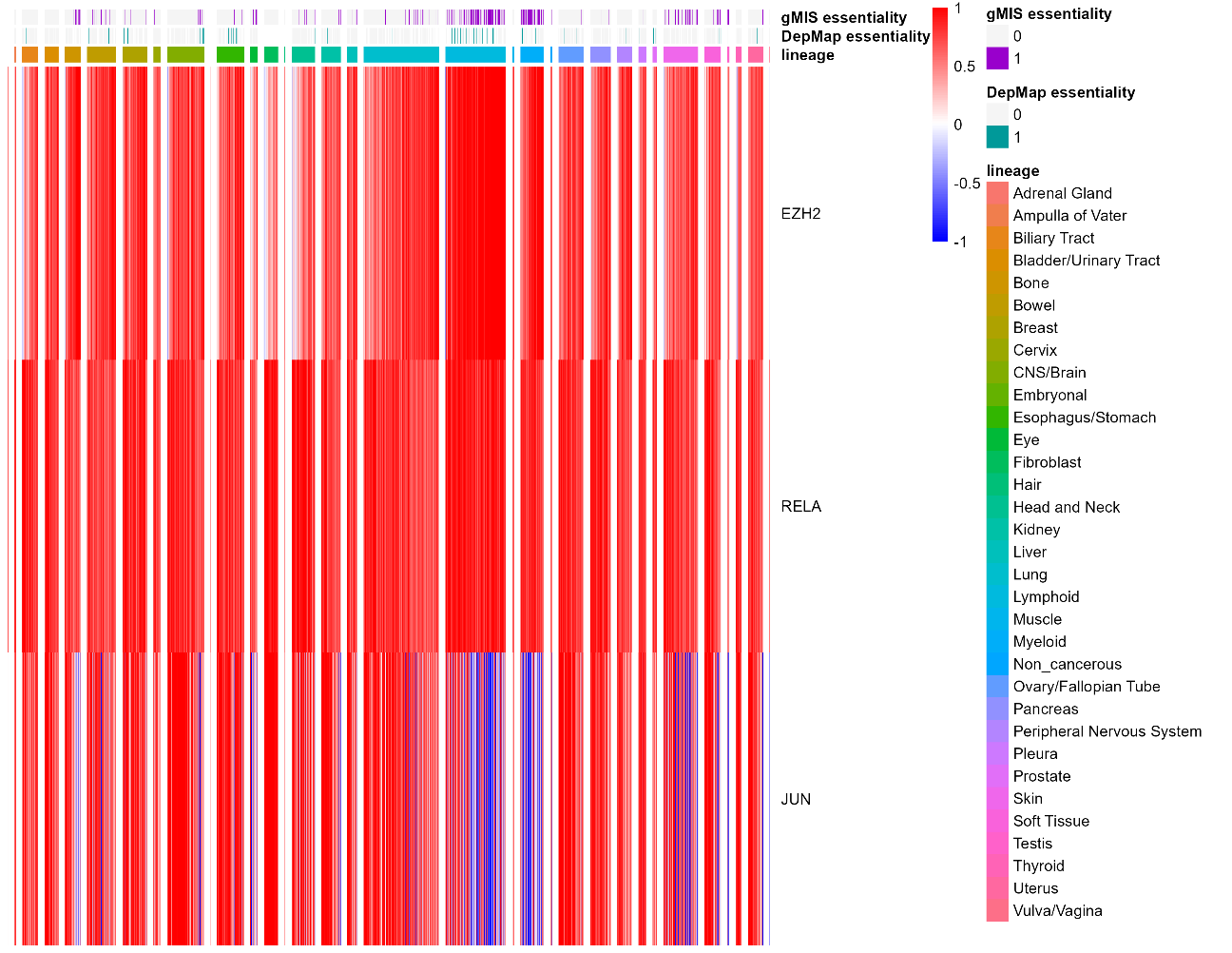

**Supplementary Figure 4. Analysis and validation of EZH2**. Heatmap of expression of the genes EZH2, RELA and JUN, which comprise a gMIS that is predicted in Human1-T2. Comparison of the essentiality of EZH2 predicted by our approach (‘*gMIS essentiality*’) and DepMap (‘*DepMap essentiality*’) for different lineages of cancer cell lines.

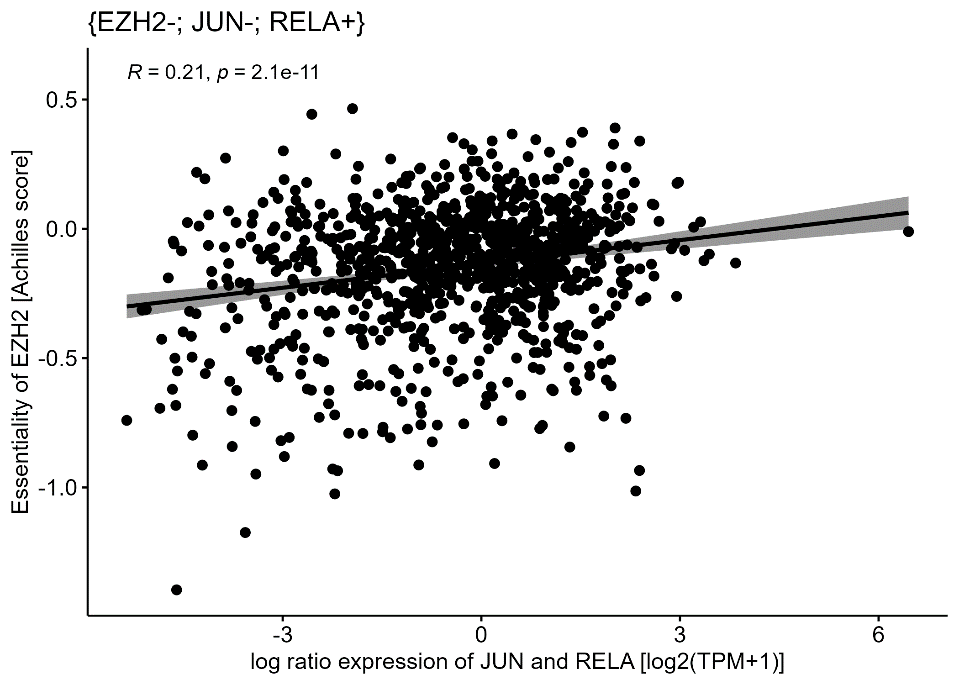

**Supplementary Figure 5.** Correlation between the essentiality of EZH2 (CRISPR knock-out screen data from DepMap) and the difference in expression between JUN and RELA in log2(TPM+1) in all DepMap cell lines.

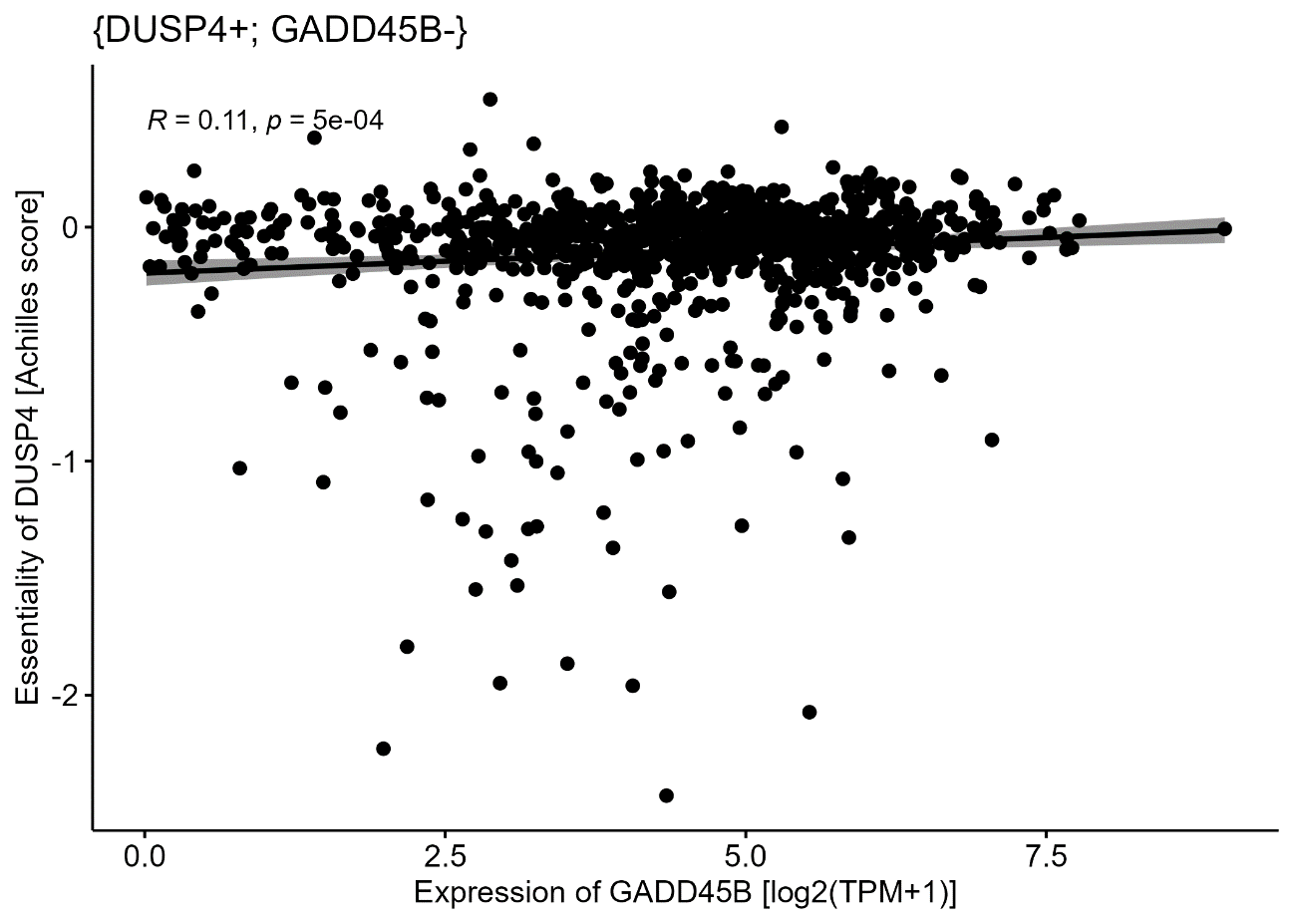

**Supplementary Figure 6.** Correlation between the essentiality of DUSP4 (CRISPR knock-out screen data from DepMap) and the expression of GADD45B in log2(TPM+1) in all DepMap cell lines.

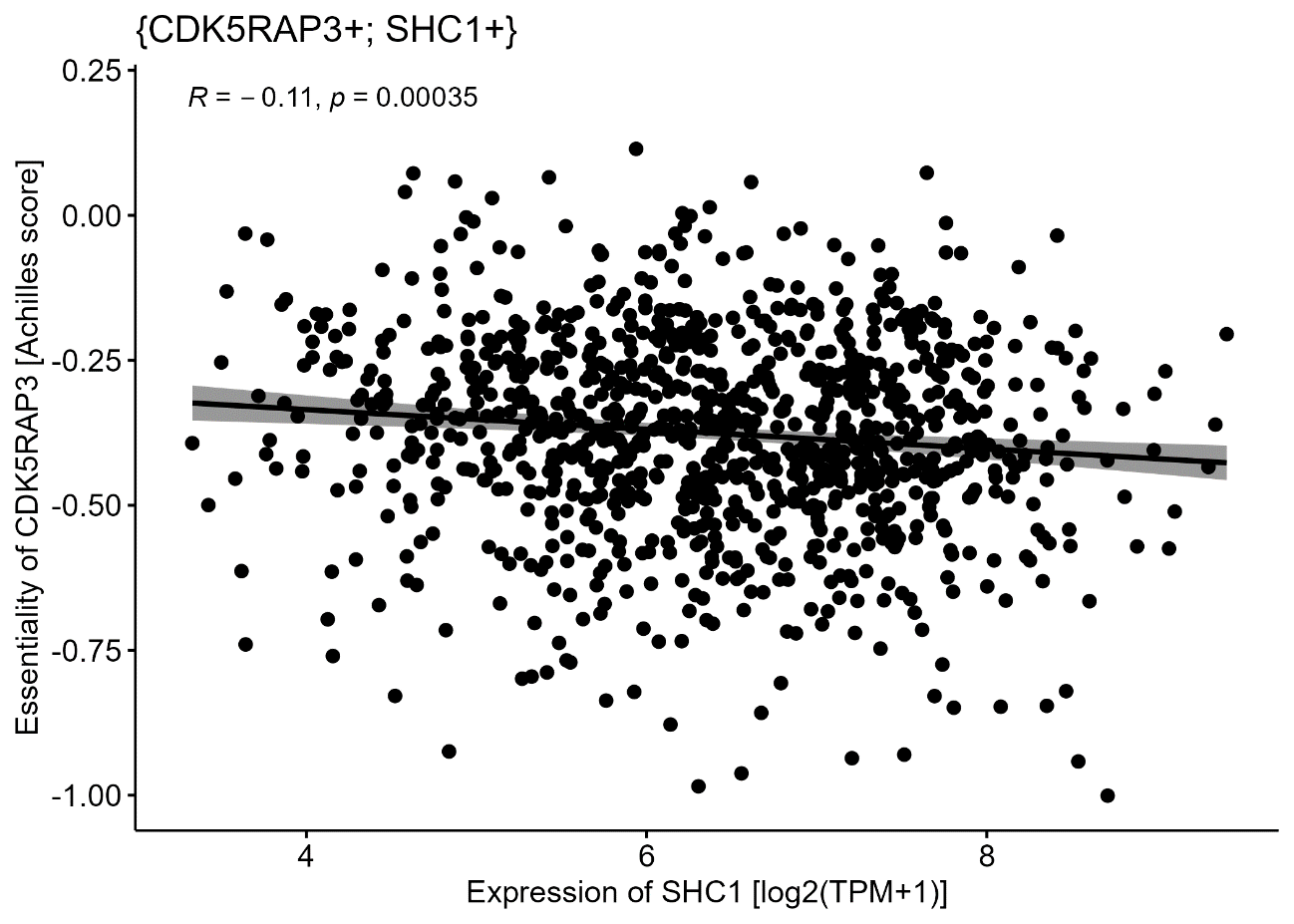

**Supplementary Figure 7.** Correlation between the essentiality of CDK5RAP3 (CRISPR knock-out screen data from DepMap) and the expression of SHC1 in log2(TPM+1) in all DepMap cell lines.

**Supplementary Tables**

**Supplementary Table 1. Description of the main features of the regulatory networks employed in the analysis.** Target metabolic genes are those reached by at least one regulatory interaction.

| **Regulatory network** | **Number of Interactions** | **Number of genes** | **Number of metabolic genes** | **Number of target metabolic genes** |
| --- | --- | --- | --- | --- |
| Omnipath | 19,964 | 6,044 | 720 | 544 |
| DoRothEA | 6,500 | 3,313 | 639 | 638 |
| TRRUST | 4,719 | 2,078 | 359 | 337 |
| Signor3.0 | 10,074 | 5,384 | 758 | 379 |

**Supplementary Table 2. Summary of multiple-layer integrated metabolic and regulatory models and computed gMCSs and gMISs***. gMCSs and gMISs until length 5 were calculated for iMR models including 1 and 2 regulatory layers. Matrix* G *were simplified to interventions involving up to 5 genes. Computation time is given in seconds (s). In the column ‘Number of Interventions (length 5)’, the value in parenthesis corresponds to the number of gMCSs/gMISs arising from the addition of the regulatory layer.*

| **Model** | **Number of genes** | **Type** | ***G* matrix**  **computation time (s)** | ***G* matrix size (length 5)** | **Number of interventions (length 5)** |
| --- | --- | --- | --- | --- | --- |
| **Human1** | 2,419 | gMCS | 483 | 1,577x11,573 | 10,091 |
| **Human1-O1** | 3,035 | gMCS | 790 | 1,800x11,573 | 14,060 (3,969) |
|  |  | gMIS | 834 | 1,928x11,573 | 14,118 (4,027) |
| **Human1-O2** | 4,274 | gMCS | 1,963 | 3,118x11,573 | 15,101 (5,010) |
|  |  | gMIS | 4,800 | 7,102x11,573 | 17,345 (7,254) |
| **Human1-D1** | 2,517 | gMCS | 1,147 | 2,024x11,573 | 11,435 (1,344) |
|  |  | gMIS | 949 | 2,146x11,573 | 11,449 (1,358) |
| **Human1-D2** | 2,527 | gMCS | 1,733 | 2,194x11,573 | 11,248 (1,157) |
|  |  | gMIS | 2,160 | 2,836x11,573 | 11,293 (1,202) |
| **Human1-T1** | 2,652 | gMCS | 790 | 1,801x11,573 | 10,779 (688) |
|  |  | gMIS | 1,040 | 1,954x11,573 | 11,695 (1,604) |
| **Human1-T2** | 2,797 | gMCS | 1,464 | 2,349x11,573 | 10,944 (853) |
|  |  | gMIS | 2,461 | 3,891x11,573 | 22,210 (12,119) |
| **Human1-S1** | 2,648 | gMCS | 670 | 1,705x11,573 | 11,820 (1,729) |
|  |  | gMIS | 768 | 1,841x11,573 | 11,829 (1,738) |
| **Human1-S2** | 3,048 | gMCS | 958 | 1,880x11,573 | 11,846 (1,755) |
|  |  | gMIS | 1,583 | 2,935x11,573 | 11,988 (1,897) |

**Supplementary Table 3.** Summary of the comparison of essentiality predictions between our gMIs approach and DepMap for different iMR models considering only synthetic lethality (SL) and both synthetic lethality and synthetic dosage lethality (SDL). Abbreviations: ‘*Human1-O1’*: integrated model with Human1 and Omnipath with one regulatory layer; ‘*Human1-O2’*: integrated model with Human1 and Omnipath with two regulatory layers; ‘*Human1-D1’*: integrated model with Human1 and DoRothEA with one regulatory layer; ‘*Human1-D2’*: integrated model with Human1 and DoRothEA with two regulatory layers; ‘*Human1-T1’*: integrated model with Human1 and TRRUST with one regulatory layer; ; ‘*Human1-T2’*: integrated model with Human1 and TRRUST with two regulatory layers; ‘*Human1-S1’*: integrated model with Human1 and Signor3.0 with one regulatory layer; ‘*Human1-S2’*: integrated model with Human1 and Signor3.0 with two regulatory layers; TP: True Positives; FP: False Positives; PPV: Positive Predicted Value; TN: True Negatives; FN: False Negatives.

| **Model** | **Type** | **TP** | **FP** | **TN** | **FN** | **PPV** | **Sensitivity** |
| --- | --- | --- | --- | --- | --- | --- | --- |
| Human1 | SL | 82.4750 | 96.5926 | 262.3536 | 28.2409 | 0.46099 | 0.7583 |
| Human1-O1 | SL | 85.7150 | 105.0725 | 279.2889 | 30.5857 | 0.44973 | 0.7489 |
|  | SDL | 86.2997 | 112.4016 | 279.9530 | 30.0078 | 0.43478 | 0.7536 |
| Human1-O2 | SL | 100.2302 | 158.4917 | 320.8482 | 37.3575 | 0.38779 | 0.7363 |
|  | SDL | 102.2449 | 163.6709 | 321.7796 | 36.3604 | 0.38488 | 0.7451 |
| Human1-D1 | SL | 84.7581 | 120.6415 | 265.1361 | 29.1263 | 0.41332 | 0.7568 |
|  | SDL | 85.0793 | 122.1401 | 266.5622 | 28.8805 | 0.41120 | 0.7591 |
| Human1-D2 | SL | 85.7777 | 123.0147 | 263.5867 | 28.2831 | 0.41145 | 0.7647 |
|  | SDL | 86.0989 | 124.5387 | 264.0304 | 27.9941 | 0.40935 | 0.7673 |
| Human1-T1 | SL | 83.0157 | 103.9843 | 274.3105 | 29.2958 | 0.44430 | 0.7521 |
|  | SDL | 83.2880 | 104.9843 | 274.1381 | 29.1959 | 0.44275 | 0.7533 |
| Human1-T2 | SL | 83.1812 | 104.1812 | 284.2194 | 30,0392 | 0.44439 | 0.7475 |
|  | SDL | 84.3800 | 106.7963 | 284.4398 | 30,0049 | 0.44176 | 0.7500 |
| Human1-S1 | SL | 86.3487 | 99.5798 | 264.4104 | 29,3232 | 0.46485 | 0.7588 |
|  | SDL | 86.5132 | 99.7150 | 265.2752 | 29,1587 | 0.46497 | 0.7601 |
| Human1-S2 | SL | 86.4437 | 99.8394 | 268.5945 | 30,0372 | 0.46448 | 0.7544 |
|  | SDL | 86.6141 | 100.2125 | 270.2106 | 29,8776 | 0.46403 | 0.7558 |

**Supplementary Table 4.** Summary list of context-specific essential genes predicted from iMR network models with TRRUST that were specifically obtained from synthetic dosage lethality interactions.

| **Gene** | **Model** | **Literature** |
| --- | --- | --- |
| ATF2 | Human1-T2 | (Liu *et al.*, 2020) |
| ATM | Human1-T2 |  |
| CPT1B | Human1-T1  Human1-T2 | (Abudurexiti *et al.*, 2020) |
| CPT1C | Human1-T2 | (Zaugg *et al.*, 2011) |
| DHODH | Human1-T1  Human1-T2 | (Zhou *et al.*, 2021)  (Boukalova *et al.*, 2020)  (Olsen *et al.*, 2022) |
| EZH2 | Human1-T2 | (Duan *et al.*, 2020)  (Kim and Roberts, 2016) |
| HDAC1 | Human1-T2 | (Yu *et al.*, 2019)  (Ropero and Esteller, 2007) |
| HDAC2 | Human1-T2 | (Jo *et al.*, 2023)  (Ropero and Esteller, 2007) |
| JUN | Human1-T2 | (Blau *et al.*, 2012)  (MacLeod *et al.*, 2019)  (Lukey *et al.*, 2016) |
| NFATC1 | Human1-T2 | (Pan *et al.*, 2013)  (Shen *et al.*, 2021) |
| PPARD | Human1-T1  Human1-T2 | (Zuo *et al.*, 2017)  (Wang *et al.*, 2016)  (Liu *et al.*, 2019) |
| RAD51 | Human1-T2 | (Z. Wang *et al.*, 2022)  (Laurini *et al.*, 2020) |
| RPIA | Human1-T1  Human1-T2 | (Chou *et al.*, 2018)  (Qiu *et al.*, 2015) |
| SUCLG1 | Human1-T1  Human1-T2 |  |
| TK1 | Human1-T1  Human1-T2 | (Bitter *et al.*, 2020)  (Jagarlamudi and Shaw, 2018)  (Topolcan and Holubec, 2008) |
| TXN | Human1-T1  Human1-T2 | (Liu *et al.*, 2023) |
| UMPS | Human1-T1  Human1-T2 |  |
| UPP1 | Human1-T2 | (Guan *et al.*, 2019)  (X. Wang *et al.*, 2022) |

**Supplementary Table 5.** Enrichment analysis with one sided hypergeometric test of our predicted tumor suppressor genes and TSGene2.0 database.

| Model | p-value | Total genes in the model | Total TSGs by TSGene | Predicted TSGs | Correct predicted TSG |
| --- | --- | --- | --- | --- | --- |
| Human1-O1 | 5.44e-02 | 3535 | 224 | 14 | 3 |
| Human1-O2 | 3.02e-11 | 4728 | 412 | 168 | 43 |
| Human1-D1 | 1.50e-02 | 3038 | 158 | 4 | 2 |
| Human1-D2 | 1.05e-11 | 3048 | 163 | 37 | 16 |
| Human1-T1 | 2.07e-03 | 3162 | 194 | 5 | 3 |
| Human1-T2 | 1.86e-06 | 3299 | 233 | 19 | 9 |
| Human1-S1 | 1 | 2663 | 131 | 2 | 0 |
| Human1-S2 | 1 | 3064 | 195 | 5 | 0 |
